# Sterically enhanced control of enzyme-assisted DNA assembly

**DOI:** 10.1101/2023.02.27.530245

**Authors:** Oliver J Irving, Lauren Matthews, Steven Coulthard, Robert K Neely, Mellissa M. Grant, Tim Albrecht

## Abstract

Traditional methods for the assembly of functionalised DNA structures, involving enzyme restriction and modification, present difficulties when working with small DNA fragments (<100bp), in part due to a lack of control over enzymatic action during the DNA modification process. This limits the design flexibility and range of accessible DNA structures. Here, we show that these limitations can be overcome by introducing chemical modifications into the DNA, which spatially restrict enzymatic activity. This approach, Sterically Controlled Nuclease Enhanced (SCoNE) DNA assembly, thereby circumvents the size limitations of conventional Gibson assembly (GA) and allows for the preparation of well-defined, functionalised DNA structures with multiple probes for specific analytes, such as IL-6, procalcitonin (PCT), and a biotin reporter group. Notably, using the same starting materials conventional GA under typical conditions fails. We demonstrate successful analyte capture based on standard and modified sandwich ELISA and also show how the inclusion of biotin probes provides additional functionality for product isolation.

## Introduction

DNA is a material with unique structural, functional, and electronic properties (1, 2). It is not only the bearer of genetic information, thereby encoding biological function, but the nucleotides as its fundamental building blocks can also provide the basis for functional structures for a range of applications in electronics, sensing, materials science via directed self-assembly (3–8). The latter concept is rooted in the work of Seeman (9) and later Rothemund et al., in the context of so-called ‘DNA origami’ (10).

A prime example of how DNA functionalisation has expanded capabilities in biomolecular sensing can be seen in resistive-pulse sensing using solid-state nanopores and nanopipettes(11–14). Originally focusing on fundamental transport and biophysical studies of (double-stranded) DNA and potentially label-free DNA sequencing (11, 12, 15–18), the introduction of functionalised DNA as a carrier for multiple capture probes for biomolecular targets has enabled both new features and applications (“carrier-enhanced” resistive-pulse sensing). Importantly, in resistive-pulse sensing the binding state of a capture probe on the carrier – target bound vs. unbound – and in principle also the concentration of the target are determined via electrical readout, with interesting prospects for device miniaturisation and point-of-care diagnostics (13, 19–21). Accordingly, the incorporation of multiple, identical capture probes positively affects the readout statistics, since multiple capture probes are detected with every DNA translocation event, ultimately reducing the total analysis time. On the other hand, DNA carriers with multiple probes for different targets enable the simultaneous detection of marker panels, which often improve detection results, compared to individual disease markers (22–25).

Unfortunately, available methods for DNA modification and functionalisation do not offer sufficient design flexibility, precision and/or scalability to produce the required structures and are typically rather labour-intensive. These are based on either the chemical or biochemical modification of already formed, double-stranded DNA or its reconstitution from single-stranded DNA with carefully chosen mismatch sequences. Specifically, chemical and biochemical modification may serve to incorporate nucleotide mismatch sequences (14, 19, 22), direct chemical (23, 24) or enzymatic modifications, in some cases followed by Click chemistry (25– 29).

The incorporation of mismatch sequences is intuitively straightforward, however in practice potentially tedious. Starting from long, double-stranded (ds) DNA, the strands are first separated into single-stranded (ss) DNA and the template strand retained. The complementary strand is replaced by a number of short ssDNA oligomers to reconstitute the dsDNA, bar in the region that is meant to be functionalised. Here, a longer piece of ssDNA with a complementary segment is used to fill the gap and to provide an overhang. The latter can then serve as a capture probe for ssDNA (30), or form the basis for further modification. Finally, enzymatic repair can be used to close any nicks along the dsDNA. However, for a long piece of ssDNA, e.g., with thousands of nucleotides, a rather large number of oligomeric units needs to be provided, potentially 100s or more. This makes the process unnecessarily complex and potentially costly (13, 19, 31).

To this end, enzymatic modification can offer a viable alternative. Enzymatic modification of DNA is a natural process which in the case of methyltransferase enzymes occurs to regulate gene expression, typically at CpG islands (32, 33). The *Thermus aquaticus* extremophile produces a methyltransferase enzyme (M.TaqI), which recognises TCGA sites, transferring the methyl group from S-Adenosyl methionine (SAM) to the adenosine residue (34). This natural process can be exploited to transfer non-native reporter, analyte capture, and/or functional groups on to DNA nucleotides and form the basis for studying methylation patterns in genomic DNA (25), cell division, or the effect of pharmaceuticals on cellular health (35–37).

A third approach to engineering DNA structures is based on DNA restriction enzymes and the use of specific restriction sites in (circular) plasmid DNA (38). Two classic examples of this are BioBrick and Golden Gate assemblies (39, 40). In BioBrick assembly, plasmids with functional genes of interest are flanked either side by specific enzyme restriction sites, referred to as BioBricks. Multiple plasmids are used for the assembly, all containing matching restriction sites. The plasmids are digested to isolate the gene fragments and the overhang regions associated with the restriction site can re-anneal, reforming a plasmid containing both genes. However, in order to create plasmids containing more genes, the complexity increases. This is due to the need for plasmids with different, orthogonal restriction sites (41), thereby increasing the probability of gene digestion, increasing the cost, and reducing the number of useable genes (42, 43). Golden gate assembly often utilises a BsaI restriction enzyme to cleave insertable DNA sequences, creating overhang regions. This occurs simultaneously with both the DNA vector and the sequences to be inserted. The overhang sequences then reform dsDNA and remaining nicks are sealed with a T4 ligase. This eliminates the restriction site, inhibiting further enzymatic action (44). Both BioBrick and Golden Gate techniques are primarily used to insert genes of interest into plasmids for microbial transformation, but typically not applied to linear DNA (45–47).

Finally, Gibson Assembly (GA) is a well-known technique for adjoining blunt ended DNA segments, without relying on enzyme restriction sites (48). It typically employs three enzymes, a 5’ exonuclease, a DNA polymerase, and a DNA ligase. The DNA segments must contain matching overlap sequences at the ends, as the 5’ exonuclease digests up to 100 bp. The now single stranded regions of the DNA overlap and combine to form dsDNA. The DNA polymerase then fills in any gaps left, and the DNA ligase then seals any remaining nicks.

However, while widely used, GA is usually inadequate for small DNA fragments, say <100 bp in length. This is because enzymatic digestion is hard to control and often leads to complete digestion of short dsDNA, at least under “standard” reaction conditions, cf. fig. 1. This therefore limits the range of accessible target designs, such as the multifunctional structures shown in the present work. In particular, our objective has been to construct DNA “carriers” with multiple capture probes for protein disease markers in well-defined locations and with potentially high density, for the use in resistive pulse sensing.

**Figure 1.**
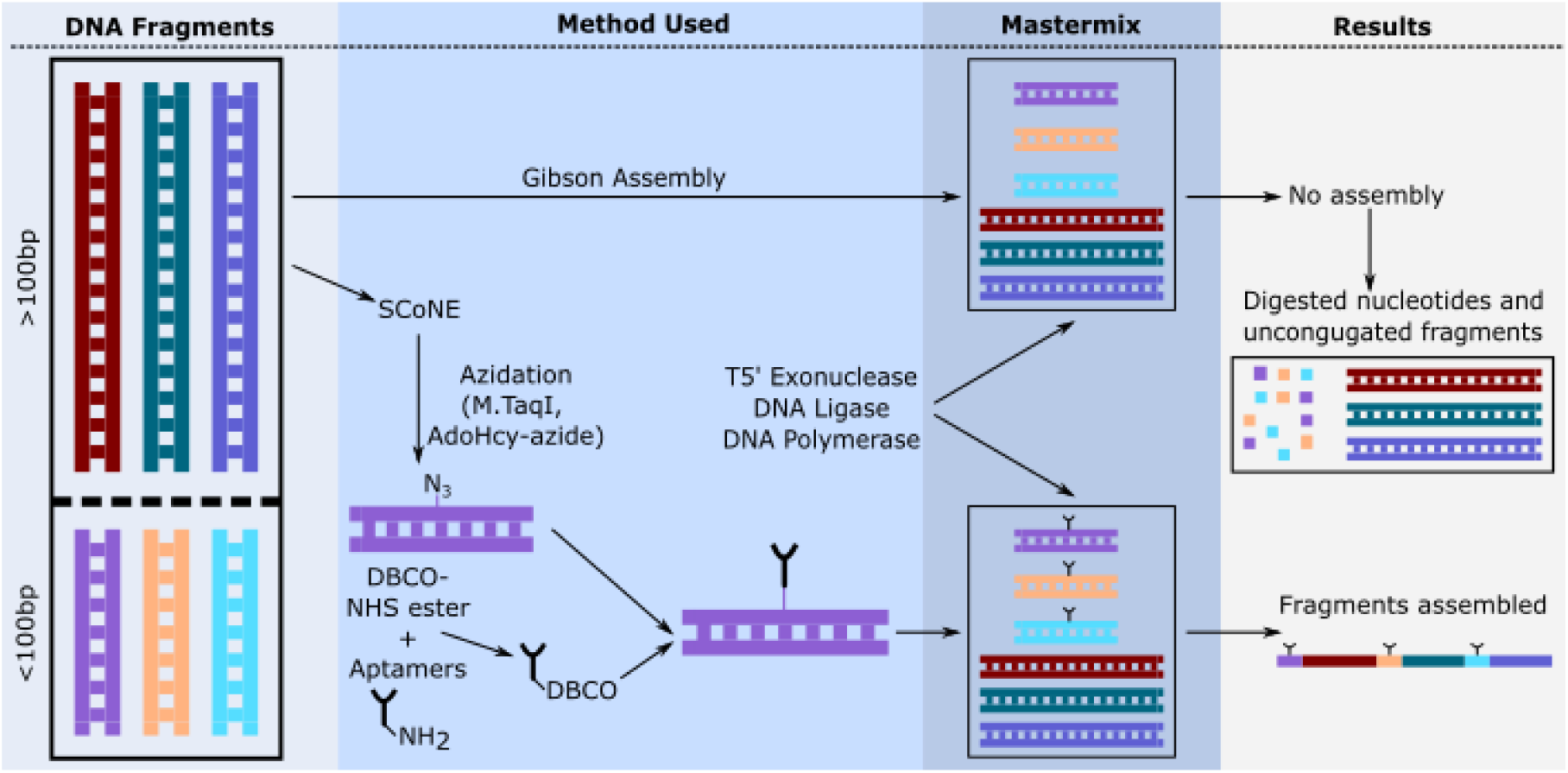
Illustration of the two preparation pathways, via conventional Gibson Assembly (top) and SCoNE assembly (bottom). The latter includes the conjugation of aminated capture probe(s) “Y” to DBCO-NHS ester and, in parallel, azidation of short (<100 bp) DNA strands using M.TaqI and AdoHcy-azide. Modified probes and the respective azidated DNA strands then undergo a copper-free Click reaction to produce ‘probe DNA’ (pDNA) for further assembly. pDNA is subsequently combined with ‘spacer DNA’ (sDNA) in an assembly mastermix to yield the final product (bottom right). In Gibson assembly, the short DNA strands remain unmodified. When the mastermix is combined with the enzymes T5’ exonuclease, Taq DNA Ligase, and Taq DNA polymerase, under standard conditions the short DNA strands are digested and cannot be assembled into the final product.

To this end, our newly developed approach, Sterically Controlled Nuclease Enhanced (SCoNE) DNA assembly, overcomes the length limitation of GA, by chemically introducing substituents into the DNA structure, which ultimately limit 5’-exonuclease activity through steric blocking of the enzyme, Figure. 2. As a result, SCoNE DNA assembly allows for the preparation of functional DNA structures, where conventional GA fails.

**Figure 2.**
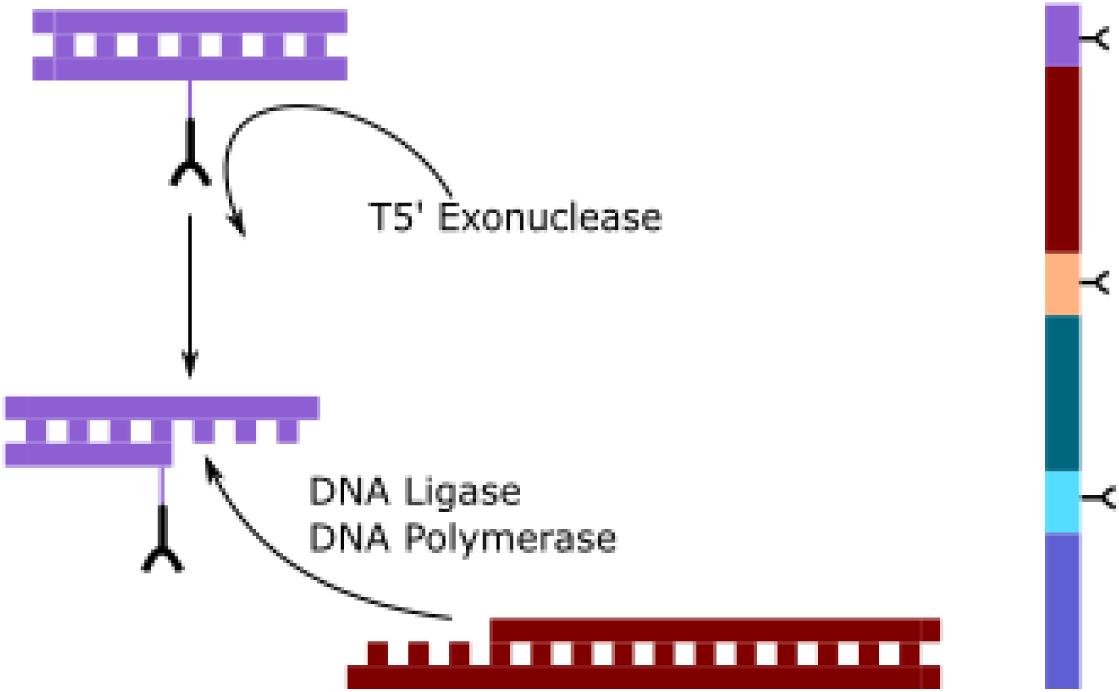
The 5’ ends of the pr- and sDNA are digested, creating complementary sticky ends. We hypothesise that the T5 exonuclease is unable to navigate across the modified probe attachment and therefore releases the DNA. Subsequently, DNA segments combine the pre-defined sequence, as illustrated in the target DNA on the right.

For SCoNE DNA assembly, we use short DNA segments (length: 34 – 100 bp), which were later modified to carry a capture probe (prDNA), and “spacer” DNA (sDNA) which in the present case is approximately 1000 bp long. As a variation of prDNA, we also prepared DNA segments containing biotin groups (iDNA), which were ultimately used to simplify isolation of the assembly product. prDNA and sDNA have a minimum 15-nucleotide overlapping region on the 3’ ends of the strand, which is complementary to the corresponding strand at their partner DNA. It is designed in such a way that prDNA and sDNA can only adjoin specifically and in the correct sequence, as shown for the target design in Figure. 1 (bottom right). While we show results for aptamer probes specific to interleukin 6 (IL-6) and PCT as well as biotin in the present work, the methodology is rather more flexible and can also be brought to incorporate other capture probes, such as antibodies (not shown).

## Methods

Details on experimental methods, DNA sequences, and conditions for Gibson and SCoNE DNA assemblies are provided in the Supplementary Information (SI), sections 1 and 2. Briefly, aminated aptamers in resuspension buffer (final concentration 10 µM, Cambio) were kept at 95°C for 5 minutes and 2 µL of this solution were added to 18 µL of DBCO-NHS ester, (50 µM, Sigma-Aldrich), incubating at room temperature on a shaker for 1.5 hours. For prDNA, 60 bp dsDNA was azidated using an AdoHcy-azide (26) and M.TaqI at 40°C for 1.5 hours. 2 µL of the aptamer-DBCO conjugate were incubated with 4 µL of the azidated dsDNA for at least 1.5 hours to form the final prDNA. To produce iDNA, the aptamer-DBCO conjugate was replaced with DBCO-dPEG®12-biotin (Sigma-Aldrich), with the remaining assembly steps remaining the same. For the final assembly step, sDNA was diluted 1:5 in nuclease-free water and added into a single reaction vial with the prDNA and iDNA, at a total DNA concentration of 1 pM. To this reaction mixture, 5 µL of nuclease-free water and 5 µL of GA mastermix (NEB) were added and incubated at 40°C for a minimum of 1.5 hours. For conventional GA, this assembly was performed with the unmodified 60 bp prDNA using 10 prDNA and 10 sDNA units. SCoNE constructs were isolated using Dynabeads™ MyOne™ Streptavidin C1 beads, via the biotin functionality on the iDNA. The concentration of the final SCoNE structures was determined by UV-Vis spectroscopy (BioSpec-nano, Shimadzu) at 260 nm, in Dynabeads™ isolation solution.

Gel electrophoresis was used to characterise the reaction products, including for sizing (1% agarose gel in 1X TAE buffer at 75 V for 45 minutes). Probe functionality was determined by ELISA and modified ELISA protocols, using trimer (2 iDNA, 1 prDNA, and 3 sDNA) and tetramer (2 iDNA, 2 prDNA, and 4 sDNA) structures.

## Results and Discussion

As a first step towards preparing a set of specific target DNA structures composed of different combinations of i-, pr- and sDNA, we compared SCoNE assembly with conventional GA. For GA, two unmodified (60 bp) prDNA and two (1 kbp) sDNA were combined in a reaction vial, as described in the Methods section above and in the SI. For SCoNE assembly, we aimed for three different designs, namely a ‘dimer’ (2 sDNA + 2 iDNA (sequences 1 and 2, see SI), total molecular weight (MW) ∼2.2 kbp), a ‘tetramer’ (4 sDNA, 2 iDNA + 2 prDNA (1 IL6 and 1 PCT aptamer on sequences 3 and 4), MW ∼4.4 kbp), and a ‘decamer’ (10 sDNA, 2 iDNA + 8 prDNA (4 IL6 and 4 PCT aptamers, sequences 3-10, MW ∼ 11 kbp). Gel electrophoresis was then used to evaluate the size of the respective DNA assembly products, as shown in Figure. 3. Lane 1 contains the 1 kbp GeneRuler; whereas lanes 2-4 were intentionally left blank to avoid intensity bleeding from the GeneRuler to the samples in lanes 5-8. Specifically, lane 5 contains the product from GA, while lanes 6-8 are SCoNE products.

**Figure 3.**
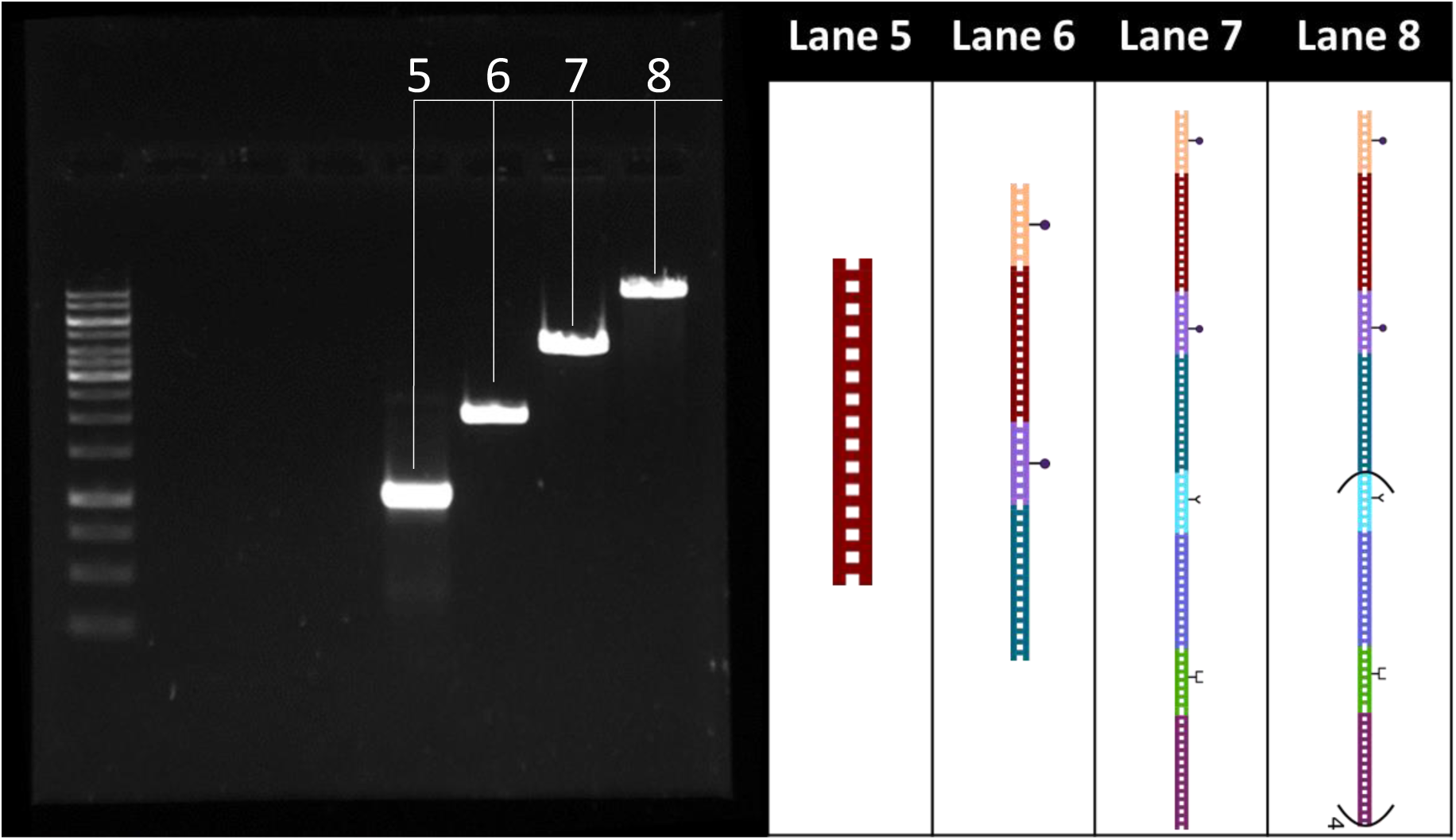
Gel electrophoresis result after Gibson and SCoNE assemblies (1% agarose gel, 75 V, 45 min.). Lane 1: 1 kbp gene ruler, lanes 2-4 blank, lane 5 Gibson assembly product and lanes 6-8 for SCoNE dimer, tetramer and decamer, respectively. Right: illustration of the hypothesized final DNA structures with biotin (circles) and aptamer (Y) probes as indicated, see main text for further discussion.

From these results, it is apparent that, in lane 5, the Gibson assembly has failed, since we observe only the template 1 kbp sDNA fragment in the gel. Conversely, for SCoNE assembly, bands consistent with correctly assembled DNA structures were found in all three cases, while those for the reactants (sDNA and pDNA) could not be detected, suggesting the reaction is highly efficient. Moreover, based on the gel results, there is no indication of significant amounts of side- or decomposition products.

For a more quantitative analysis, the migration distance was measured from the centre of the initial well to the centre of the DNA band in each case. The bands for the DNA ladder were used to produce a calibration curve and a mono-exponential least-square fit performed (y = A1*exp(-x/t1) + y0 (49, 50)), Figure. 4. Based on this model, the actual weight of the respective SCoNE and sDNA structures were interpolated, assuming that the capture probes themselves had a negligible effect on DNA migration. The latter assumption seems justified, given their small relative contribution to the charge and mass of the constructs. From this analysis, the DNA sizes were estimated to be 900 ± 200 bp for sDNA (expected ∼1 kbp), 2000 ± 200 bp (expected ∼2.2 kbp), for the dimer, 4100 ± 200 bp (expected ∼4.4 kbp), for the tetramer, and 11100 ± 500 bp (expected ∼11 kbp) for the decamer structure. Hence, the correspondence between the actual and expected values is very good, which is further supported by the correlation plot in Figure. 4, inset (Gradient: 1.01 ± 0.05). We provide further details on our efforts to optimise the reaction conditions and confirm some of our findings in the SI section 3 but will now proceed to demonstrate the functionality of the prepared DNA constructs.

**Figure 4.**
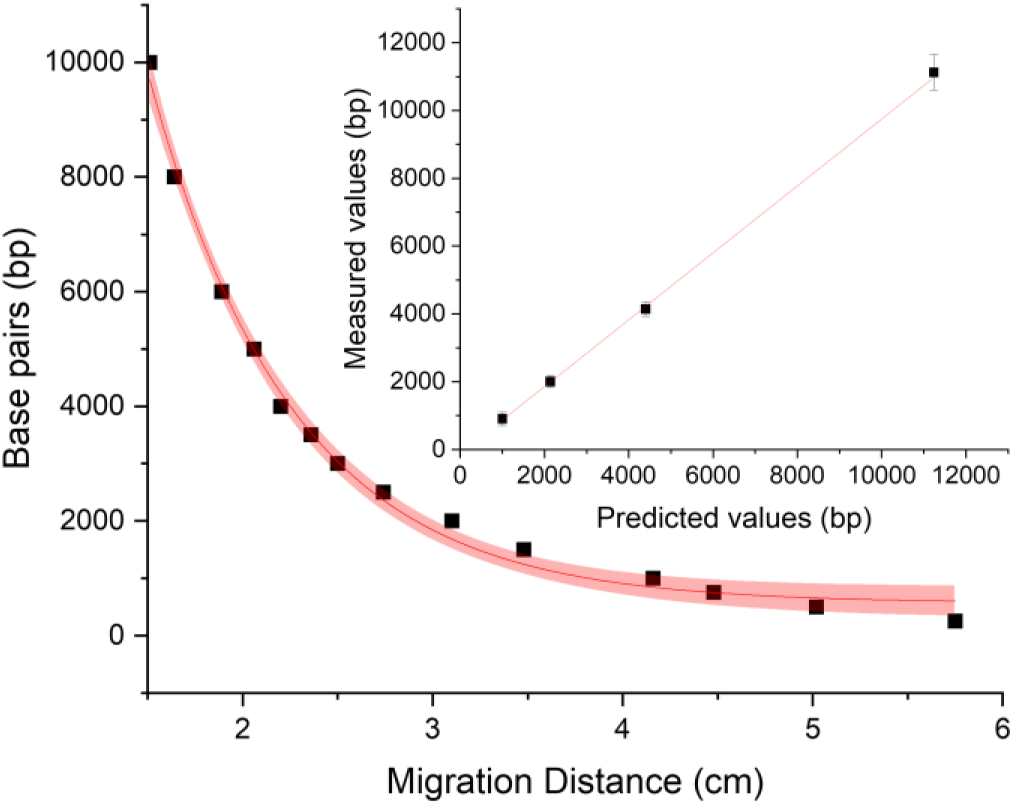
Calibration curve calculated from the 1kbp GeneRuler ladder with a mono-exponential fit (red, solid line; shaded: 95% confidence interval), and the resultant sDNA and SCoNE fragment size calculated from this model. Inset: correlation plot (Gradient: 1.01 ± 0.05).

In order to demonstrate the functionality of the prepared DNA constructs, we developed sandwich ELISA protocols that incorporate the SCoNE assemblies. We benchmarked this against a standard ELISA to assess the ability of the DNA structure to bind a single, specific analyte (protocol 1, timer). Modified protocols were employed to determine whether targets were bound simultaneously, for example by detecting the presence of functional biotin groups via Avidin Binding Complex (ABC) and direct Horseradish Peroxidase (HRP) quantification (protocol 2, trimer); by using (bound) IL6 for capture and PCT for quantification (or the reverse) (protocol 3a and b respectively, tetramer); and in order to validate the specificity of target binding via ABC/HRP detection (protocol 4, tetramer). Further details can be found in section 2 of the SI.

From UVvis spectroscopy measurements, the SCoNE DNA concentrations were found to be 16±1 ng/µL for protocol 1 and for protocol 2 with IL6 as primary and PCT as secondary antibodies; 8±1 ng/µL for protocol 2 with PCT as primary and IL6 as secondary antibodies and protocol 3; 10±1 ng/µL for protocol 4 with IL6 as primary and PCT as secondary antibodies. Final protein yields were compared to these values.

The results are shown in Figure. 5. Following protocol 1, 443±65 pg/mL of protein was captured, corresponding to a capture efficiency of (73.7±10.8)%, whereas in protocol 2, 419±23 pg/mL was captured, equivalent to a capture efficiency of (69.7±3.8)%. For protocol 3a with PCT as primary and IL-6 as secondary antibodies, 126±33 pg/mL was captured with a capture efficiency of (44.0±11.6)%, whereas in the inverse configuration (protocol 3b), i.e. with IL-6 as primary and PCT as secondary antibody, 103±19 pg/mL or (28.4±5.2)% was captured. In all of these cases, the signal output demonstrates that each capture is functional and specific to the respective target protein.

**Figure 5.**
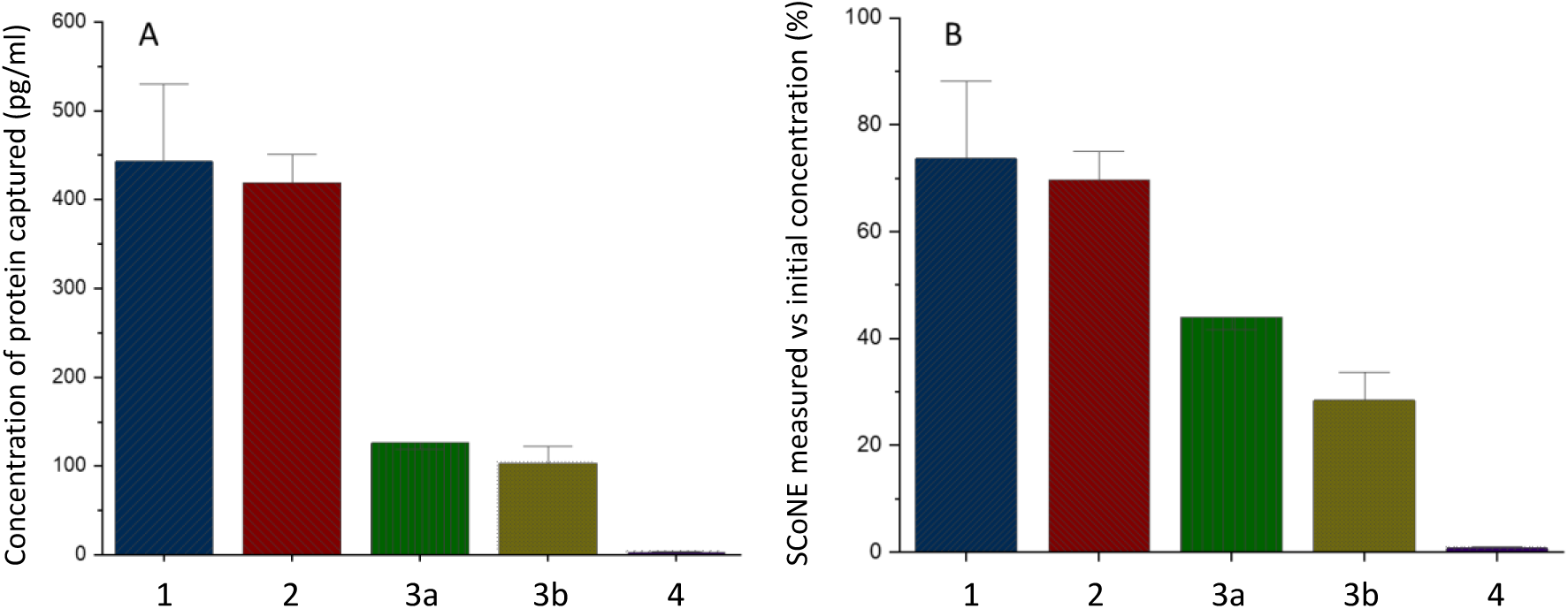
A) ELISA results comparing the concentration of target protein captured by the respective SCoNE structures, and B) the corresponding capture efficiency (UVvis measurements, N=6), where ‘1’ is a standard sandwich ELISA for IL-6 using a SCoNE trimer, ‘2’ a sandwich ELISA for PCT (secondary antibody removed, biotin groups used as a measure for protein concentration), ‘3’ a sandwich ELISA for both PCT (primary antibody) and IL-6 (secondary antibody) using a SCoNE tetramer, ‘4’ is a sandwich ELISA for both IL-6 (primary antibody) and PCT (secondary antibody) using a SCoNE tetramer, and ‘5’ a sandwich ELISA for PCT (primary antibody) using an IL-6 secondary antibody without incubation with IL-6.

On the contrary, for protocol 4, a low protein concentration of 2.8±0.6 pg/mL was measured, corresponding to a capture efficiency of only (0.8±0.2)%. Due to the low signal generated when no secondary protein is present, this therefore indicates that biotin sites were blocked efficiently, further supporting the validity of the results obtained from protocols 1-3 (by excluding non-specific binding).

## Conclusion

We have introduced a method for the assembly of functionalised DNA, which overcomes some limitations of conventional Gibson Assembly. Specifically, it allows for the assembly of short DNA segments below approximately 100 bp by sterically restricting nuclease activity to the respective end regions of the DNA. As a proof-of-concept, we have prepared several DNA designs, with well-defined sizes ranging from 2.2 to 11 kbp and interchangeable conjugate groups. The modularity of the approach and the ability to attach a range of different capture probe for biomolecular targets implies significant flexibility, in terms of target design and application. Here, we have focused on IL6- and PCT-specific aptamers as capture probes, in a repeat or alternating configuration, and incorporated biotin groups to simplify the isolation of the final construct.

To test the functionality of the SCoNE constructs, we performed standard and modified ELISA protocols. The efficiency of trimer structures for capturing protein was determined to be 73.7±10.8% while the tetramer had 44.0±11.6 and 28.4±5.2% for protocol 3a and 3b respectively. The ELISA results also showed that the biotin group could be used directly for sensing and blocked effectively when not required.

Whilst we have focused on the conjugation of aptamers here, we note that the assembly method could be applied to other amine containing probe or reporter groups without requiring significant protocol alteration. To this end, initial experiments with IL-6 and PCT-specific antibodies have proven successful, highlighting the design flexibility of the approach.

## Availability of data and materials

Data related to this work is publicly available via the University of Birmingham’s UBIRA e-respository at DOI [to be supplied] and upon request from the authors.

## Authors’ contribution

TA conceived the project. OJI conceived and developed the technique, tested the protocol and optimised the construction parameters. LM tested the technique and provided additional data. RKN advised on the development of methylase-directed DNA modification and SC synthesised and tested the cofactor analogues used for DNA modification. MG advised on biochemical aspects, including binding assays. All authors contributed to the writing of the manuscript.

## Acknowledgements

Authors wish to thank Anton Vladyka and Christopher Weaver for their assistance in developing data extraction, and handling MATLAB code used in this work.

## Notes

### Competing Interest Statement

The authors have declared no competing interest.

