## Supplemental Information for "Sterically enhanced control of enzyme-assisted DNA assembly"

Oliver J Irving[1], Lauren Matthews[1], Steven Coulthard[1], Robert K Neely[1], Mellissa M. Grant[2], and Tim Albrecht[1]*

[1]School of Chemistry, University of Birmingham, Edgbaston, Birmingham B15 2TT, UK

[2]School of Dentistry, Institute of Clinical Sciences, University of Birmingham and Birmingham Dental Hospital (Birmingham Community Healthcare Trust), 5 Mill Pool Way, Edgbaston, Birmingham B5 7EG, UK

*

### 1. DNA Segment sequences

The following sequences were used for the spacer DNA (sDNA) and probe DNA (prDNA) strands:

| prDNA | Sequence (5'-3') |
|---|---|
| 1 | CTGCAACGCAGCGCTTGTAGGATTCACATCGAACGGATTCAGTGCTTATCGCATAGACAG |
| 2 | GAACGGCCCGGTGTGCTTGTGATTCTAGTCGACGCGATTCAACGTTTGGAAATCCCATAC |
| 3 | GTATTGAGGGGTGTGCCACTGATTCTGATCGAAACGATTCTAGCGCACGACCTGGCAGAG |
| 4 | GATCAATGCTCCAACGAACTGATTCATCTCGAGGCGATTCAAAGCGGGAATGATACGGCG |
| 5 | AACCACTATAACTAGAGCAGGATTCACATCGAAGCGATTCACGTGTTGGCACCGGATATG |
| 6 | CTGTAGCAGCCTGAGTAGGTGATTCAATTCGATATGATTCGCGCTCTCTAATTTGCGTCG |
| 7 | TGCCAGTCTTGGCCTCTTCAGATTCGTGTCGACCAGATTCATTCGTGCATTACGTTATCG |
| 8 | TCGGCTGACTAGAGAATAGGGATTCTTGTCGACCCGATTCCCCCGCTTCTGCTTGAACAC |
| 9 | CGGACAAAGAGGCAGCGAATGATTCATGTCGACCCGATTCGGTTAGGACCGTCAGTTATG |
| 10 | ATTGACACTATTAGTCCAGAGATTCAATTCGATAAGATTCCGCAGGACCGTGCTCGTAGA |

| sDNA | Sequence (5'-3') |
|---|---|
| 1 | AGTGCTTATCGCATAGACAGCGAGGACACTTGGCACGACCAAGCATACTTGTGAGCGGAAGTCTCGCAATCCTTCGTGCCAGGACCTGGCGGCCCGTGCTCGCTGATAGCCG<br>TTAAAATGGTTTTGGGGTTGTTCATCGGACACTATCCCACTACTATTGCATCACGTACACTACACCCCTGGGGAGTAATCGGGACTGAAACCCACTTCGTGAAGATATGCGAGAT<br>CTCATAAACGTACGGCATGGTACACGACTATATTCATGCCCCGCCGTCATATGAAAGGGGAACATTATGAAAACAATGTGTAGTATCCCGCCACCTACCTGTCTTTTCAGGCTAA<br>GGCGCGCGACACAGGATTGGGAAGGATGCCCATATGTGTGCTCATCTCAAACAACCTTAACCCATTAGCCAGGTTACAGCAGAGCGCATAACAACGACGGACCAGCATGATCT<br>TCCAACGAACACTCTCTTGCAACGACACGTTACAAATAACGTGCGTCACGCTGTTACATGTTTTCTAGCATCAGCGGGCTTGAGACCTGAAGCAACCTTGTATCTAGGATCTCGC<br>CGGAAATCTACGAACCTCACCCGACGATGATACTTTGTTGCCCAGAGCGAGGACACTCAATAGACCTTTCAGACTGTAGAATTATCACACCAAAGAGTCAGCACGGTTTCGCGA<br>GAAATGCACACGGCGGTCGCTCAATCCATGGTATAATTATTCACTGCTCAGCGTAGGGTCATGAACTGTGGCGCTTGTGGCTCCCCGTGAAGTACAATGGGAACCGGCAACGG<br>TCTAGCATGGGAATTTCTTTACTCTGCAATCGCGAGTATGAGAAAGTCGGCAACGCAGCCAGTGTAAAATACCCCTTTCACGGGAATTCCGGAACACAGCTCGCGGTGGCTAAA<br>TCGCCCTAAGGGAAGGGTGTTCCAACTTCAAATTTGTAAATTTTCTTAATTGTATATAAAGTTTCCGGTGAACGGCCCGGTGTGCTTGT |
| 2 | AACGTTTGGAAATCCCATACCGAGGACACTTGGCACGACCAAGCATACTTGTGAGCGGAAGTCTCGCAATCCTTCGTGCCAGGACCTGGCGGCCCGTGCTCGCTGATAGCCGT<br>TAAAATGGTTTTGGGGTTGTTCATCGGACACTATCCCACTACTATTGCATCACGTACACTACACCCCTGGGGAGTAATCGGGACTGAAACCCACTTCGTGAAGATATGCGAGATC<br>TCATAAACGTACGGCATGGTACACGACTATATTCATGCCCCGCCGTCATATGAAAGGGGAACATTATGAAAACAATGTGTAGTATCCCGCCACCTACCTGTCTTTTCAGGCTAAG<br>GCGCGCGACACAGGATTGGGAAGGATGCCCATATGTGTGCTCATCTCAAACAACCTTAACCCATTAGCCAGGTTACAGCAGAGCGCATAACAACGACGGACCAGCATGATCTT<br>CCAACGAACACTCTCTTGCAACGACACGTTACAAATAACGTGCGTCACGCTGTTACATGTTTTCTAGCATCAGCGGGCTTGAGACCTGAAGCAACCTTGTATCTAGGATCTCGCC<br>GGAAATCTACGAACCTCACCCGACGATGATACTTTGTTGCCCAGAGCGAGGACACTCAATAGACCTTTCAGACTGTAGAATTATCACACCAAAGAGTCAGCACGGTTTCGCGAG<br>AAATGCACACGGCGGTCGCTTAATCCATGGTATAATTATTCACTGCTCAGCGTAGGGTCATGAACTGTGGCGCTTGTGGCTCCCCGTGAAGTACAATGGGAACCGGCAACGGT<br>CTAGCATGGGAATTTCTTTACTCTGCAATCGCGAGTATGAGAAAGTCGGCAACGCAGCCAGTGTAAAATACCCCTTTCACGGGAATTCCGGAACACAGCTCGCGGTGGCTAAAT<br>CGCCCTAAGGGAAGGGTGTTCCAACTTCAAATTTGTAAATTTTCTTAATTGTATATAAAGTTTCCGGTGTATTGAGGGGTGTGCCACT |

| | |
|---|---|
| **3** | TAGCGCACGACCTGGCAGAGCGAGGACACTTGGCACGACCAAGCATACTTGTGAGCGGAAGTCTCGCAATCCTTCGTGCCAGGACCTGGCGGCCCGTGCTCGCTGATAGCCG<br>TTAAAATGGTTTTGGGGTTGTTCATCGGACACTATCCCACTACTATTGCATCACGTACACTACACCCCTGGGGAGTAATCGGGACTGAAACCCACTTCGTGAAGATATGCGAGAT<br>CTCATAAACGTACGGCATGGTACACGACTATATTCATGCCCCGCCGTCATATGAAAGGGGAACATTATGAAAACAATGTGTAGTATCCCGCCACCTACCTGTCTTTTCAGGCTAA<br>GGCGCGCGACACAGGATTGGGAAGGATGCCCATATGTGTGCTCATCTCAAACAACCTTAACCCATTAGCCAGGTTACAGCAGAGCGCATAACAACGACGGACCAGCATGATCT<br>TCCAACGAACACTCTCTTGCAACGACACGTTACAAATAACGTGCGTCACGCTGTTACATGTTTTCTAGCATCAGCGGGCTTGAGACCTGAAGCAACCTTGTATCTAGGATCTCGC<br>CGGAAATCTACGAACCTCACCCGACGATGATACTTTGTTGCCCAGAGCGAGGACACTCAATAGACCTTTCAGACTGTAGAATTATCACACCAAAGAGTCAGCACGGTTTCGCGA<br>GAAATGCACACGGCGGTCGCTTAATCCATGGTATAATTATTCACTGCTCAGCGTAGGGTCATGAACTGTGGCGCTTGTGGCTCCCCGTGAAGTACAATGGAACCGGCAACGG<br>TCTAGCATGGGAATTTCTTTACTCTGCAATCGCGAGTATGAGAAAGTCGGCAACGCAGCCAGTGTAAAATACCCCTTTCACGGGAATTCCGGAACACAGCTCGCGGTGGCTAAA<br>TCGCCCTAAGGGAAGGGTGTTCCAACTTCAAATTTGTAAATTTTCTTAATTGTATATAAAGTTTCCGGTGATCAATGCTCCAACGAACT |
| **4** | AAAGCGGGAATGATACGGCGCGAGGACACTTGGCACGACCAAGCATACTTGTGAGCGGAAGTCTCGCAATCCTTCGTGCCAGGACCTGGCGGCCCGTGCTCGCTGATAGCCG<br>TTAAAATGGTTTTGGGGTTGTTCATCGGACACTATCCCACTACTATTGCATCACGTACACTACACCCCTGGGGAGTAATCGGGACTGAAACCCACTTCGTGAAGATATGCGAGAT<br>CTCATAAACGTACGGCATGGTACACGACTATATTCATGCCCCGCCGTCATATGAAAGGGGAACATTATGAAAACAATGTGTAGTATCCCGCCACCTACCTGTCTTTTCAGGCTAA<br>GGCGCGCGACACAGGATTGGGAAGGATGCCCATATGTGTGCTCATCTCAAACAACCTTAACCCATTAGCCAGGTTACAGCAGAGCGCATAACAACGACGGACCAGCATGATCT<br>TCCAACGAACACTCTCTTGCAACGACACGTTACAAATAACGTGCGTCACGCTGTTACATGTTTTCTAGCATCAGCGGGCTTGAGACCTGAAGCAACCTTGTATCTAGGATCTCGC<br>CGGAAATCTACGAACCTCACCCGACGATGATACTTTGTTGCCCAGAGCGAGGACACTCAATAGACCTTTCAGACTGTAGAATTATCACACCAAAGAGTCAGCACGGTTTCGCGA<br>GAAATGCACACGGCGGTCGCTTAATCCATGGTATAATTATTCACTGCTCAGCGTAGGGTCATGAACTGTGGCGCTTGTGGCTCCCCGTGAAGTACAATGGAACCGGCAACGG<br>TCTAGCATGGGAATTTCTTTACTCTGCAATCGCGAGTATGAGAAAGTCGGCAACGCAGCCAGTGTAAAATACCCCTTTCACGGGAATTCCGGAACACAGCTCGCGGTGGCTAAA<br>TCGCCCTAAGGGAAGGGTGTTCCAACTTCAAATTTGTAAATTTTCTTAATTGTATATAAAGTTTCCGGTAACCACTATAACTAGAGCAG |
| **5** | ACGTGTTGGCACCGGATATGCGAGGACACTTGGCACGACCAAGCATACTTGTGAGCGGAAGTCTCGCAATCCTTCGTGCCAGGACCTGGCGGCCCGTGCTCGCTGATAGCCG<br>TTAAAATGGTTTTGGGGTTGTTCATCGGACACTATCCCACTACTATTGCATCACGTACACTACACCCCTGGGGAGTAATCGGGACTGAAACCCACTTCGTGAAGATATGCGAGAT<br>CTCATAAACGTACGGCATGGTACACGACTATATTCATGCCCCGCCGTCATATGAAAGGGGAACATTATGAAAACAATGTGTAGTATCCCGCCACCTACCTGTCTTTTCAGGCTAA<br>GGCGCGCGACACAGGATTGGGAAGGATGCCCATATGTGTGCTCATCTCAAACAACCTTAACCCATTAGCCAGGTTACAGCAGAGCGCATAACAACGACGGACCAGCATGATCT<br>TCCAACGAACACTCTCTTGCAACGACACGTTACAAATAACGTGCGTCACGCTGTTACATGTTTTCTAGCATCAGCGGGCTTGAGACCTGAAGCAACCTTGTATCTAGGATCTCGC<br>CGGAAATCTACGAACCTCACCCGACGATGATACTTTGTTGCCCAGAGCGAGGACACTCAATAGACCTTTCAGACTGTAGAATTATCACACCAAAGAGTCAGCACGGTTTCGCGA<br>GAAATGCACACGGCGGTCGCTTAATCCATGGTATAATTATTCACTGCTCAGCGTAGGGTCATGAACTGTGGCGCTTGTGGCTCCCCGTGAAGTACAATGGAACCGGCAACGG<br>TCTAGCATGGGAATTTCTTTACTCTGCAATCGCGAGTATGAGAAAGTCGGCAACGCAGCCAGTGTAAAATACCCCTTTCACGGGAATTCCGGAACACAGCTCGCGGTGGCTAAA<br>TCGCCCTAAGGGAAGGGTGTTCCAACTTCAAATTTGTAAATTTTCTTAATTGTATATAAAGTTTCCGGTCTGTAGCAGCCTGAGTAGGT |
| **6** | GCGCTCTCTAATTTGCGTCGCGAGGACACTTGGCACGACCAAGCATACTTGTGAGCGGAAGTCTCGCAATCCTTCGTGCCAGGACCTGGCGGCCCGTGCTCGCTGATAGCCG<br>TTAAAATGGTTTTGGGGTTGTTCATCGGACACTATCCCACTACTATTGCATCACGTACACTACACCCCTGGGGAGTAATCGGGACTGAAACCCACTTCGTGAAGATATGCGAGAT<br>CTCATAAACGTACGGCATGGTACACGACTATATTCATGCCCCGCCGTCATATGAAAGGGGAACATTATGAAAACAATGTGTAGTATCCCGCCACCTACCTGTCTTTTCAGGCTAA<br>GGCGCGCGACACAGGATTGGGAAGGATGCCCATATGTGTGCTCATCTCAAACAACCTTAACCCATTAGCCAGGTTACAGCAGAGCGCATAACAACGACGGACCAGCATGATCT<br>TCCAACGAACACTCTCTTGCAACGACACGTTACAAATAACGTGCGTCACGCTGTTACATGTTTTCTAGCATCAGCGGGCTTGAGACCTGAAGCAACCTTGTATCTAGGATCTCGC<br>CGGAAATCTACGAACCTCACCCGACGATGATACTTTGTTGCCCAGAGCGAGGACACTCAATAGACCTTTCAGACTGTAGAATTATCACACCAAAGAGTCAGCACGGTTTCGCGA<br>GAAATGCACACGGCGGTCGCTTAATCCATGGTATAATTATTCACTGCTCAGCGTAGGGTCATGAACTGTGGCGCTTGTGGCTCCCCGTGAAGTACAATGGAACCGGCAACGG<br>TCTAGCATGGGAATTTCTTTACTCTGCAATCGCGAGTATGAGAAAGTCGGCAACGCAGCCAGTGTAAAATACCCCTTTCACGGGAATTCCGGAACACAGCTCGCGGTGGCTAAA<br>TCGCCCTAAGGGAAGGGTGTTCCAACTTCAAATTTGTAAATTTTCTTAATTGTATATAAAGTTTCCGGTTGCCAGTCTTGGCCTCTTCA |
| **7** | ATTCGTGCATTACGTTATCGCGAGGACACTTGGCACGACCAAGCATACTTGTGAGCGGAAGTCTCGCAATCCTTCGTGCCAGGACCTGGCGGCCCGTGCTCGCTGATAGCCGT<br>TAAAATGGTTTTGGGGTTGTTCATCGGACACTATCCCACTACTATTGCATCACGTACACTACACCCCTGGGGAGTAATCGGGACTGAAACCCACTTCGTGAAGATATGCGAGATC<br>TCATAAACGTACGGCATGGTACACGACTATATTCATGCCCCGCCGTCATATGAAAGGGGAACATTATGAAAACAATGTGTAGTATCCCGCCACCTACCTGTCTTTTCAGGCTAAG<br>GCGCGCGACACAGGATTGGGAAGGATGCCCATATGTGTGCTCATCTCAAACAACCTTAACCCATTAGCCAGGTTACAGCAGAGCGCATAACAACGACGGACCAGCATGATCTT<br>CCAACGAACACTCTCTTGCAACGACACGTTACAAATAACGTGCGTCACGCTGTTACATGTTTTCTAGCATCAGCGGGCTTGAGACCTGAAGCAACCTTGTATCTAGGATCTCGCC<br>GGAAATCTACGAACCTCACCCGACGATGATACTTTGTTGCCCAGAGCGAGGACACTCAATAGACCTTTCAGACTGTAGAATTATCACACCAAAGAGTCAGCACGGTTTCGCGAG<br>AAATGCACACGGCGGTCGCTTAATCCATGGTATAATTATTCACTGCTCAGCGTAGGGTCATGAACTGTGGCGCTTGTGGCTCCCCGTGAAGTACAATGGAACCGGCAACGGT<br>CTAGCATGGGAATTTCTTTACTCTGCAATCGCGAGTATGAGAAAGTCGGCAACGCAGCCAGTGTAAAATACCCCTTTCACGGGAATTCCGGAACACAGCTCGCGGTGGCTAAAT<br>CGCCCTAAGGGAAGGGTGTTCCAACTTCAAATTTGTAAATTTTCTTAATTGTATATAAAGTTTCCGGTTCGGCTGACTAGAGAATAGG |
| **8** | CCCCGCTTCTGCTTGAACACCGAGGACACTTGGCACGACCAAGCATACTTGTGAGCGGAAGTCTCGCAATCCTTCGTGCCAGGACCTGGCGGCCCGTGCTCGCTGATAGCCG<br>TTAAAATGGTTTTGGGGTTGTTCATCGGACACTATCCCACTACTATTGCATCACGTACACTACACCCCTGGGGAGTAATCGGGACTGAAACCCACTTCGTGAAGATATGCGAGAT<br>CTCATAAACGTACGGCATGGTACACGACTATATTCATGCCCCGCCGTCATATGAAAGGGGAACATTATGAAAACAATGTGTAGTATCCCGCCACCTACCTGTCTTTTCAGGCTAA<br>GGCGCGCGACACAGGATTGGGAAGGATGCCCATATGTGTGCTCATCTCAAACAACCTTAACCCATTAGCCAGGTTACAGCAGAGCGCATAACAACGACGGACCAGCATGATCT<br>TCCAACGAACACTCTCTTGCAACGACACGTTACAAATAACGTGCGTCACGCTGTTACATGTTTTCTAGCATCAGCGGGCTTGAGACCTGAAGCAACCTTGTATCTAGGATCTCGC<br>CGGAAATCTACGAACCTCACCCGACGATGATACTTTGTTGCCCAGAGCGAGGACACTCAATAGACCTTTCAGACTGTAGAATTATCACACCAAAGAGTCAGCACGGTTTCGCGA<br>GAAATGCACACGGCGGTCGCTTAATCCATGGTATAATTATTCACTGCTCAGCGTAGGGTCATGAACTGTGGCGCTTGTGGCTCCCCGTGAAGTACAATGGAACCGGCAACGG<br>TCTAGCATGGGAATTTCTTTACTCTGCAATCGCGAGTATGAGAAAGTCGGCAACGCAGCCAGTGTAAAATACCCCTTTCACGGGAATTCCGGAACACAGCTCGCGGTGGCTAAA<br>TCGCCCTAAGGGAAGGGTGTTCCAACTTCAAATTTGTAAATTTTCTTAATTGTATATAAAGTTTCCGGTCGGACAAAGAGGCAGCGAAT |
| **9** | GGTTAGGACCGTCAGTTATGCGAGGACACTTGGCACGACCAAGCATACTTGTGAGCGGAAGTCTCGCAATCCTTCGTGCCAGGACCTGGCGGCCCGTGCTCGCTGATAGCCG<br>TTAAAATGGTTTTGGGGTTGTTCATCGGACACTATCCCACTACTATTGCATCACGTACACTACACCCCTGGGGAGTAATCGGGACTGAAACCCACTTCGTGAAGATATGCGAGAT<br>CTCATAAACGTACGGCATGGTACACGACTATATTCATGCCCCGCCGTCATATGAAAGGGGAACATTATGAAAACAATGTGTAGTATCCCGCCACCTACCTGTCTTTTCAGGCTAA<br>GGCGCGCGACACAGGATTGGGAAGGATGCCCATATGTGTGCTCATCTCAAACAACCTTAACCCATTAGCCAGGTTACAGCAGAGCGCATAACAACGACGGACCAGCATGATCT<br>TCCAACGAACACTCTCTTGCAACGACACGTTACAAATAACGTGCGTCACGCTGTTACATGTTTTCTAGCATCAGCGGGCTTGAGACCTGAAGCAACCTTGTATCTAGGATCTCGC<br>CGGAAATCTACGAACCTCACCCGACGATGATACTTTGTTGCCCAGAGCGAGGACACTCAATAGACCTTTCAGACTGTAGAATTATCACACCAAAGAGTCAGCACGGTTTCGCGA<br>GAAATGCACACGGCGGTCGCTTAATCCATGGTATAATTATTCACTGCTCAGCGTAGGGTCATGAACTGTGGCGCTTGTGGCTCCCCGTGAAGTACAATGGAACCGGCAACGG<br>TCTAGCATGGGAATTTCTTTACTCTGCAATCGCGAGTATGAGAAAGTCGGCAACGCAGCCAGTGTAAAATACCCCTTTCACGGGAATTCCGGAACACAGCTCGCGGTGGCTAAA<br>TCGCCCTAAGGGAAGGGTGTTCCAACTTCAAATTTGTAAATTTTCTTAATTGTATATAAAGTTTCCGGTATTGACACTATTAGTCCAGA |
| **10** | CGCAGGACCGTGCTCGTAGACGAGGACACTTGGCACGACCAAGCATACTTGTGAGCGGAAGTCTCGCAATCCTTCGTGCCAGGACCTGGCGGCCCGTGCTCGCTGATAGCCG<br>TTAAAATGGTTTTGGGGTTGTTCATCGGACACTATCCCACTACTATTGCATCACGTACACTACACCCCTGGGGAGTAATCGGGACTGAAACCCACTTCGTGAAGATATGCGAGAT<br>CTCATAAACGTACGGCATGGTACACGACTATATTCATGCCCCGCCGTCATATGAAAGGGGAACATTATGAAAACAATGTGTAGTATCCCGCCACCTACCTGTCTTTTCAGGCTAA<br>GGCGCGCGACACAGGATTGGGAAGGATGCCCATATGTGTGCTCATCTCAAACAACCTTAACCCATTAGCCAGGTTACAGCAGAGCGCATAACAACGACGGACCAGCATGATCT<br>TCCAACGAACACTCTCTTGCAACGACACGTTACAAATAACGTGCGTCACGCTGTTACATGTTTTCTAGCATCAGCGGGCTTGAGACCTGAAGCAACCTTGTATCTAGGATCTCGC<br>CGGAAATCTACGAACCTCACCCGACGATGATACTTTGTTGCCCAGAGCGAGGACACTCAATAGACCTTTCAGACTGTAGAATTATCACACCAAAGAGTCAGCACGGTTTCGCGA<br>GAAATGCACACGGCGGTCGCTTAATCCATGGTATAATTATTCACTGCTCAGCGTAGGGTCATGAACTGTGGCGCTTGTGGCTCCCCGTGAAGTACAATGGAACCGGCAACGG<br>TCTAGCATGGGAATTTCTTTACTCTGCAATCGCGAGTATGAGAAAGTCGGCAACGCAGCCAGTGTAAAATACCCCTTTCACGGGAATTCCGGAACACAGCTCGCGGTGGCTAAA<br>TCGCCCTAAGGGAAGGGTGTTCCAACTTCAAATTTGTAAATTTTCTTAATTGTATATAAAGTTTCCGGTAATTCGTCGGACTGCTGGAT |

### 2. SCoNE methodology

To create a SCoNE structure, the technique is split into several independent stages. All reactions and incubations were performed in low DNA binding PCR tubes (ThermoFisher) and dilutions using nuclease-free Water for Molecular Biology (Merck) unless otherwise stated. All general purifications were performed using a PCR clean-up kit (GenElute$^{TM}$, Sigma-Aldrich) using 20 µL of nuclease-free water for elution, unless stated. First, the wash solution was prepared by adding 12 mL of the wash solution concentrate to 48 mL of 100% ethanol (HPLC grade, Sigma-Aldrich). The mini spin column was inserted into the provided collection tube.

To the column, 0.5 mL of the column preparation solution was added and centrifuged at 13,500 x g for 1 minute. For this procedure, all centrifugation steps were performed at 13,500 x g. The solution was discarded, and the column re-inserted into the collection tube. To the sample, 5 times its volume of binding buffer was added and mixed well. The solution was then transferred into the column and centrifuged for 1 minute. The elute was then discarded and the column re-inserted into the collection tube. 0.5 mL of the diluted wash solution was then added to the column and centrifuged. The elute was then discarded, the column re-inserted into the collection tube and centrifuged for 2 minutes. The column was transferred to a new collection tube and incubated with 20 µL of nuclease-free water (Merck) for 1 minute. This was then centrifuged for 1 minute, the elute collected in a separate collection tube, and stored at -20°C until use. All enzymes stated were acquired from NEB. DNA concentration determination was performed using 2 µL of sample and UV-Vis spectroscopy, measuring the absorbance at 260 nm (BioSpec-nano,Shimadzu).

2.1. Preparation of sDNA

sDNA was obtained as a section of a plasmid form from GeneArt (Invitrogen), flanked by SfiI enzyme restriction sites. The sequences were generated at random (SI section 1) with designed regions on both 5' and 3' ends to match corresponding prDNA segments. The code to generate these sequences is available by request. SfiI was used to excise the 1kbp sDNA segment from the whole plasmid using the conditions provided in Table 1. Each sDNA segment was excised independently.

Table S1. Conditions for SfiI digestion of sDNA segments

| Reagent | Volume (µL) |
|---|---|
| SfiI(20,000 units/mL) | 1 |
| Plasmid (600ng/µL) | 2 |
| 10X Cutsmart buffer | 5 |
| Nuclease free water | 42 |

The reaction was performed at 50°C for 1 hour then purified using a prior to PCR amplification. This was performed to isolate the 1 kbp sDNA segment, increasing the efficiency of the PCR reaction, and ensuring only the 1 kbp segment of interest was amplified. Concentration was determined using UV-visible spectroscopy. A 25 µL PCR reaction was performed using a Taq PCR Kit (NEB in a 0.5 mL PCR tube). The volumes of reaction buffers and components were added as described in Table 2. Due to differing yields of DNA from the SfiI digestion, the volume of DNA and nuclease free water was adjusted to compensate for this difference.

Table S2. Component and volume requirements for PCR

| Component | Volume to add in a 25µL reaction (µL) | Final concentration |
|---|---|---|
| 10X standard Taq rection buffer | 2.5 | 1X |
| 10mM dNTPs | 0.5 | 200 µM |
| 10µM forward primer | 0.5 | 0.2 µM |
| 10µM reverse primer | 0.5 | 0.2 µM |
| Template DNA | 0.8-1.2 | 10 nM |
| Taq DNA polymerase | 0.125 | 1.25 enzymatic units |
| Nuclease free water | 19.675 -20.075 | / |

This solution was gently homogenised using a wide pipette, to prevent breaking of the DNA fragments, before being gently spun down in a microcentrifuge. The PCR tube was placed in a thermocycler (PrimeG, Techne) under the conditions described in Table 3. Where rows are highlighted in light grey, these steps were cycled together in the order written in the table.

Table S3. PCR cycle protocol

| Cycle step | Temperature (°C) | Time (s) | Cycles performed |
|---|---|---|---|
| Denaturation (initial) | 95 | 30 | 1 |
| Denaturation | 95 | 20 | 30 |
| Annealing | 50 | 30 | |
| Extension | 68 | 60 | |
| Extension (final) | 72 | 300 | 1 |
| Hold | 4 | Until ready to remove | |

Final DNA concentration was determined using UV-vis spectroscopy (BioSpec-nano, Shimadzu) at 260 nm, and additional PCR reactions performed until a minimum final concentration of 600 ng/µL was achieved.

2.2 Probe preparation

Aminated aptamers (5’NH2, Cambio) were briefly centrifuged and resuspended in Resuspension Buffer (#RTW0001, Cambio) using 11.2 µL and 24.9 µL for the human procalcitonin aptamer and the human IL-6 aptamer, respectively, for a 100X working concentration (ATW0060-GM3-25 and ATW0035-GM3-25 respectively, Cambio). Aptamers were selected based on commercial availability and high affinity (26.6 nM and 19 nM respectively). These were aliquoted and stored at -20°C until use.

To prepare dibenzocyclooctyne (DBCO) –NHS ester, 1 mg of DBCO-NHS-ester (Sigma-Aldrich) was added to 1 mL of HPLC grade Dimethyl sulfoxide (DMSO, Sigma-Aldrich) to create a 2.5 mM stock. To 49 µL of nuclease free water (Merck), 1 µL of the 2.5 mM DBCO solution was added to create a working stock of 50 µM.

Aptamers were diluted to a 10X working concentration (10 µM) in aptamer folding buffer (RTW0003, Cambio), and heated to 95°C for 5 minutes, then left to cool to room temperature for 15 minutes prior to use. 2 µL of the prepared aptamer solution was added into a new PCR tube and diluted with 18 µL of a 50 µM DBCO-NHS ester solution and homogenised well. This was incubated at room temperature for a minimum of 1.5 hours on a shaker to allow conjugation of the probe to the DBCO.

Probes such as antibodies, affimers, and fluorescent tags, containing a free amine group can be used in place of the aptamer, however this is currently under investigation.

2.3 prDNA azidation

Each prDNA was prepared in a separate vial under the same conditions. The reaction vessel contained 1 µL of the prDNA (IDT) strand at 600 ng/µL, 2 µL Cutsmart buffer 10X, 1 µL AdoHcy-azide donor at 100 µM (illustration of the molecule as shown in Figure S1), 0.5 µL M.TaqI (NEB) and 15.5 µL of nuclease free water (Merck), adding the enzyme last. Each vial was incubated at 40°C for 1.5 hours. After which, 0.5 µL proteinase K (800 units/mL, NEB) was added and incubated at 40°C for 1 hour.

Vials were removed from the heat and allowed to cool to room temperature for 20 minutes. The solution was then purified using a PCR clean-up kit (GenElute™, Sigma-Aldrich), and stored at 4°C until required.

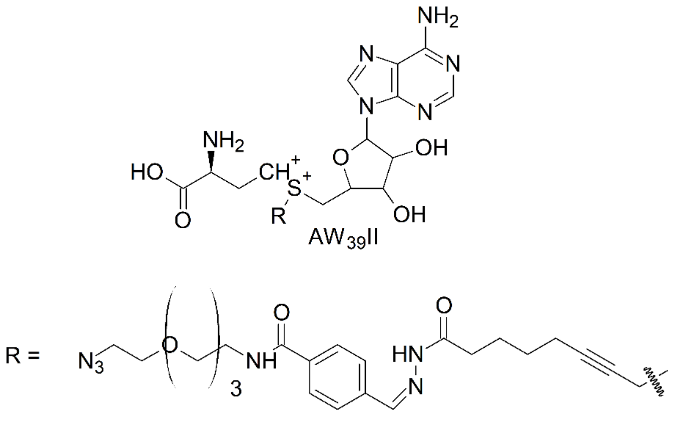


Figure S1. Illustration of AdoHcy-azide, with the SAM molecule modified with the azide containing R group as opposed to the standard methyl group.

2.4 prDNA and iDNA assembly and conjugation

Assembly of the probe strands was performed in separate vials. To each reaction vial, 2 µL of the desired probe solution was added to 4 µL of the azidated prDNA strand. This was incubated at room temperature on a shaker for a minimum of 1.5 hours to allow the click reaction to take place. To prepare iDNA strands, the same method was used replacing the prepared aptamer with 2 µL of 50 µM DBCO-dPEG®12-biotin (Sigma-Aldrich). The vials were stored at room temperature for up to 2 days prior to use.

2.5 Final assembly

All final assembly steps were performed in the same vial. sDNA segments were diluted 1:5 in nuclease-free water to create a 1X working solution. sDNA segments and prDNA segments were added into a single vial at equal concentrations (1:1 ratio), to achieve a final 1 pM solution, and homogenised well using a wide pipette tip. 10 µL of this solution was then removed and pipetted into a separate vial. 5 µL of a Gibson assembly master mix and 5 µL of nuclease free water added and homogenised gently. Vials were then incubated at 40°C for a minimum of 1.5 hours. SCoNE structures were assembled in a variety of different sizes, from dimers (2 prDNA or iDNA and 2 sDNA segments) to decamers (10 prDNA or iDNA and 10 sDNA segments).

2.6 Gel electrophoresis

A 1% agarose gel was used for all fragment separation and analysis. To 100 mL of 1X Tris-acetate-EDTA buffer (TAE, Tris (hydroxymethyl) aminomethane 1g, EDTA, tetrasodium < 1g, Water 98 mL, Disodium EDTA ~1g, 1,3-Propanediol, 2-amino-2-(hydroxymethyl)-, acetate (salt) ~1g, Fisher Scientific), 1 g of agarose powder (Sigma-Aldrich) was added. This was microwaved for 2 minutes until the agarose had completely dissolved. The solution was allowed to cool to 50°C then poured into a gel tray with an 8 well comb in place and incubated at room temperature for 30 minutes for the gel to set. To 5 µL of each DNA sample, and DNA ladder (GeneRuler 1 kbp DNA Ladder, ThermoFisher), 1 µL of 6X DNA loading buffer (ThermoFisher) was added and homogenised. The gel was then placed into a Mini-Sub Cell GT Cell (BIORAD), and this filled with 1X TAE buffer until the gel was covered. The well comb was then removed and the first well filled with 5 µL of a DNA ladder (GeneRuler 1 kbp DNA Ladder, ThermoFisher). The remaining wells were filled with 5 µL of the DNA samples under analysis or DNA standards. After connecting the unit to the power pack, the gel was run at 75 V for 45 minutes. Once finished, the gel was then placed in 1X GelRed (Sigma-Aldrich) for 45 minutes before imaging in a UV illumination box (BIORAD). The gel images were then analysed using Fiji-ImageJ.

2.7 SCoNE extraction using Streptavidin beads

SCoNE structures were extracted using Dynabeads™ MyOne™ Streptavidin C1 beads (ThermoFisher). Three assembly reaction vials were combined, and contents were added to 70 µL of Dynabeads™. This was incubated at room temperature on a shaker for 10 minutes. This was then placed in a magnetic vial holder and incubated on the benchtop for 3 minutes. The vial was removed from the magnetic holder, the supernatant discarded, and beads washed with 200 µL of 80% EtOH, and placed back on the shaker. The wash step was repeated 3 times. Following the third wash, the vial was opened, and the contents dried in air for 5 minutes. To elute the SCoNE DNA, 50 µL of nuclease free water was added and incubated on the shaker for 10 minutes. The vial was then incubated on the magnetic holder for 3 minutes and the supernatant collected and stored at -20°C prior to use.

2.8 ELISA

To determine the ability of the probe groups binding analytes of interest, standard, and modified ELISA experiments were used and developed, as illustrated in Figure S2.7. ELISA experiments were split into four separate categories: standard sandwich ELISA, direct biotin binding ELISA, counter protein ELISA, and missing protein ELISA. For the first, third, and fourth ELISAs, streptavidin (Sigma-Aldrich) was added in 3X excess to bind biotin extraction groups, to prevent nonspecific measurements. Assembled SCoNE structures were incubated at room temperature with human IL-6 (Sigma-Aldrich), procalcitonin (Sigma-Aldrich), or both at 50 ng/µL for 20 minutes on a shaker. SCoNE structures were then re-isolated using Dynabeads™ as previously described, with 300 µL of nuclease free water (Merck) used for isolation. This was done to create enough volume to cover the ELISA well plate base. For non-streptavidin blocking experiments, SCoNE structures could not be isolated using this technique, therefore additional washing steps were added to the ELISA protocol. For IL-6 and procalcitonin experiments, ELISA kits used were obtained from Stratech (orb390920-BOR-96Tests) and Merck (RAB0037-1KT) respectively.

Protein standards were set up in a serial dilution from 300 pg/mL to 4.69 pg/mL using the sample dilution solution, provided in each ELISA kit, as the blank. To each well of a precoated 96 well plate, 100 µL of protein standard and/or isolated SCoNE structures were added, the wells sealed, and incubated at 37°C for 90 minutes. The cover was then removed, the supernatant discarded, the wells blotted onto paper towels, and then washed with 0.01 M phosphate buffer solution (PBS, pH 7.4) three times. For non-streptavidin blocking experiments, the plate was washed 5 times. After each wash step the wells were blotted onto paper towels. To each well, 100 µL of the biotinylated secondary antibody was added and incubated at 37°C for 60 minutes. The supernatant discarding and wash step was then repeated before addition of 100µL of an avidin-biotin-peroxidase (ABC) solution to each well and incubated at 37°C for 30 minutes. The wells were then washed using 0.01 M PBS five times, allowing the PBS to sit in the wells for 2 minutes on each wash prior to blotting on paper towels. To each well, 90 µL of TMB solution was then added and incubated at 37°C for 20 minutes. After incubation, 100 µL of TMB stop solution was added to each well, and the absorbance read at 450 nm on a microplate reader (Infinite 200 PRO microplate reader, Tecan Trading AG, Switzerland).

To measure the ability of SCoNE structures to bind analytes of interest, modifications were made to standard ELISA methods. To measure a trimer SCoNE structure with its biotin groups blocked using streptavidin, a standard ELISA was used (ELISA A). Here SCoNE bound protein was bound to a primary IL-6 antibody (2.A.1). Secondary IL-6 antibody was then bound to the protein (2.A.2). ABC was then bound to the biotin labelled antibody (2.A.3), and TMB converted to a coloured compound for absorbance measurement (2.A.4). To measure the direct interaction with the biotin groups on the SCoNE structure, a direct biotin binding ELISA was developed, using a trimer SCoNE structure (2.B). The SCoNE bound protein was bound to a primary IL-6 antibody (2.B.1), but the secondary antibody was removed from the experimental procedure (2.B.2). ABC was then bound to the biotin labelled antibody (2.B.3), and TMB converted to a coloured compound for absorbance measurement (2.B.4). A modified ELISA using the tetramer SCoNE structure, where the primary antibody binds procalcitonin and the secondary binds IL-6, was developed as a counter protein ELISA (2.C). This was done to measure the ability of the SCoNE structures to capture multiple analytes. SCoNE bound protein was bound to a primary procalcitonin antibody (2.C.1). A secondary IL-6 antibody was added, binding to the IL-6 protein (2.C.2). ABC was then bound to the biotin labelled IL-6 antibody (2.C.3), and TMB converted to a coloured compound for absorbance measurement (2.C.4). This was also performed using the reverse antibody configuration where the primary antibody captured IL-6 and the secondary measured procalcitonin. Finally, to determine blocking of the biotin groups using streptavidin, specificity of the aptamer, and specificity of the secondary antibody, a missing protein ELISA was developed (ELISA D). A SCoNE structure was incubated with procalcitonin only, leaving the IL-6 aptamer unbound. SCoNE bound protein was bound to primary procalcitonin antibody (2.D.1). A secondary IL-6 antibody was added but was unable to bind (2.D.2). Due to the lack of free biotin groups, the ABC cannot bind (2.D.3), and therefore with the addition of TMB, it was hypothesised that no colour change would occur (2.D.4). As protein concentration was determined through use of a single aptamer bound to a SCoNE structure, the concentration of SCoNE structures was calculated based on a 1:1 molarity ratio. This value was converted to pg/mL using the total molecular mass of the structure (2080103 g/mol for a trimer and 2090503 g/mol for a tetramer). This was then converted to ng/µL, the dilution factor applied, and the final values compared against those obtained from UV-vis spectroscopy measurements.

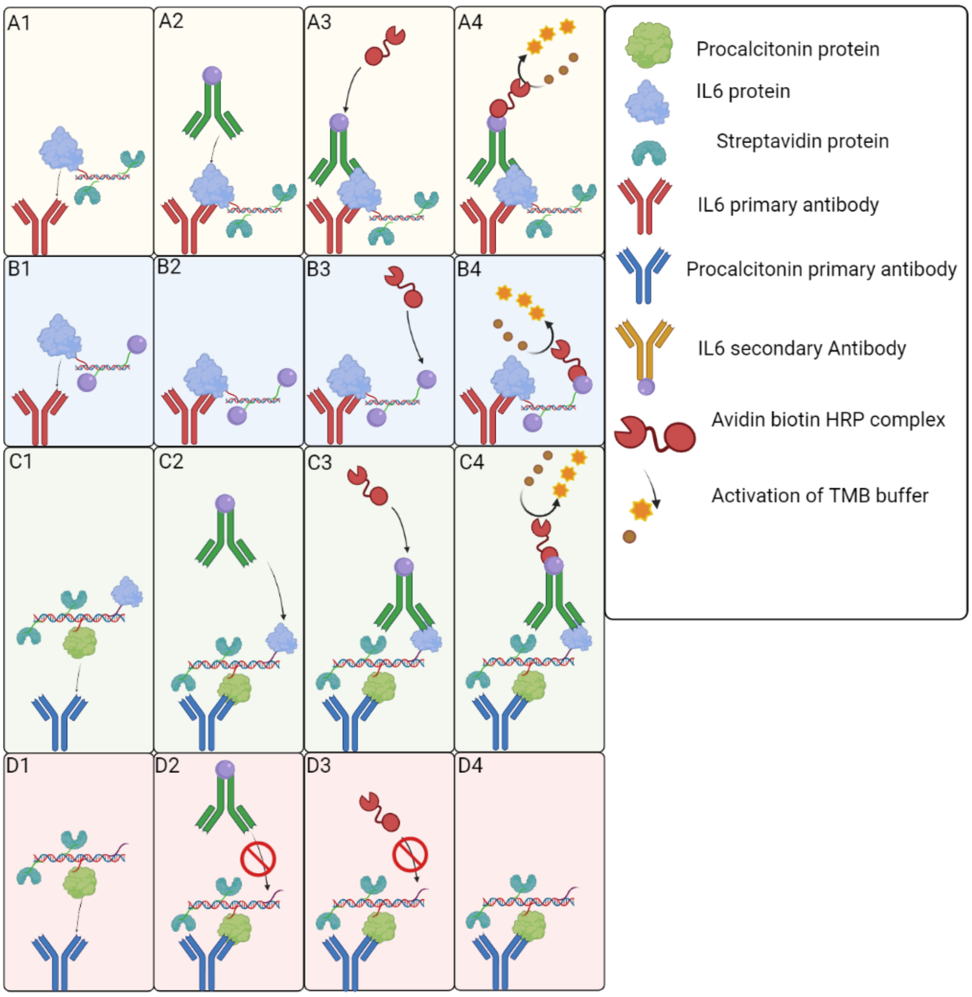


Figure S2. Standard and modified sandwich ELISA protocols. A) Typical sandwich ELISA using the trimer SCoNE structure for protein isolation using streptavidin to bind reactive biotin groups. A1: SCoNE bound protein binding to primary IL6 antibody; A2: secondary IL-6 antibody binding to protein; A3: ABC binding to biotin labelled antibody; A4: ABC converting TMB buffer to coloured compound for measurement. B) Modified sandwich ELISA using the trimer SCoNE structure, but removing the secondary antibody and measuring the protein concentration based on the biotin binding alone. B1: SCoNE bound protein binding to primary IL6 antibody; B2: Secondary IL-6 antibody not added; B3: ABC binding to biotin labelled SCoNE; B4: ABC converting TMB buffer to coloured compound for measurement. C) Modified sandwich ELISA using the tetramer SCoNE structure where the primary antibody binds procalcitonin and the secondary binds IL-6. C1: SCoNE bound protein binding to primary procalcitonin antibody; C2: secondary IL-6 antibody binding to IL6 protein; C3: ABC binding to biotin labelled IL6 antibody; C4: ABC converting TMB buffer to coloured compound for measurement; D) Similar to C using the tetramer SCoNE structure without incubation with IL6: secondary antibody cannot bind and no signal observable. D1: SCoNE bound protein binding to primary procalcitonin antibody; D2: secondary IL6 antibody added but cannot bind anything so is washed off; D3: ABC cannot bind as biotin sites are blocked by streptavidin D4: ABC is not present so cannot convert TMB to colour.

### 3 Optimisation of SCoNE assembly

To this end, a range of protocol parameters were modified to improve the assembly and yield of the final product, as summarised in table 4. This included the initial combined DNA concentration during the assembly reaction, which was varied between 0.125 and 5 pM. Despite our best efforts and for reasons that are not entirely clear at this point, the formation of SCoNE constructs was not observed below 0.5 pM or above 1 pM, providing a rather narrow concentration range where the reaction completed successfully. Secondly, we explored whether DMSO, required for DBCO NHS ester preparation but with a potentially negative impact on aptamer function, could be avoided by using water-soluble sulfo-DBCO instead. However, we found that when using the sulfo-DBCO NHS ester for prDNA assembly, the DNA yield was reduced. We hypothesise that the presence of the negatively charged sulfonate group inhibits interaction with the DNA, decreasing the conjugation efficiency (49). Finally, Gibson assembly is recommended to be performed at 50°C, denaturing the 5' exonuclease quickly. However, we found that lower yields were obtained when using the recommended temperature, compared to our optimised values. The inclusion of the biotin linker in place of the aptamer was shown not to impact the assembly reaction and to ng/µL, the dilution factor applied, and the final values compared against those obtained from UV-vis spectroscopy measurements.

Table S4. Optimisation parameters explored for the development of SCoNE methodology with outcomes confirmed by gel electrophoresis.

| Optimisation parameter | Condition applied | Observations |
|---|---|---|
| Starting DNA concentration for assembly reaction | 0.125, 0.25, 0.5, 1, 5 pM | SCoNE structure formation was not observed below 0.5 pM or above 1 pM. No significant difference in gray values obtained from the gel was observed between 0.5 and 1 pM starting concentration. |
| DBCO-NHS ester | DBCO and sulfo-DBCO NHS ester | Higher concentration of prDNA assembled using DBCO-NHS ester . |
| Temperature during enzymatic assembly step | 30, 40, 45, 50, 60°C | No assembly observed below 30°C or above 50 or. Low yield of formation at 45°C and 50°C (<50% of that achieved at 40°C). Best yield observed at 40°C. |

From the results of this optimisation we have been able to greatly improve the initial protocol to develop a robust method develop to reliably synthesise DNA nanostructures of desired lengths.
